# Stabilizing microtubules aids neurite structure and disrupts syncytia formation in human cytomegalovirus-infected human forebrain neurons

**DOI:** 10.1101/2024.08.16.608340

**Authors:** Jacob W Adelman, Andrew T Sukowaty, Kaitlyn J Partridge, Jessica E. Gawrys, Scott S. Terhune, Allison D. Ebert

## Abstract

Human cytomegalovirus (HCMV) is a prolific human herpesvirus that infects most individuals by adulthood. While typically asymptomatic in adults, congenital infection can induce serious neurological symptoms including hearing loss, visual deficits, cognitive impairment, and microcephaly in 10-15% of cases. HCMV has been shown to infect most neural cells with our group recently demonstrating this capacity in stem cell-derived forebrain neurons. Infection of neurons induces deleterious effects on calcium dynamics and electrophysiological function paired with gross restructuring of neuronal morphology. Here, we utilize an iPSC-derived model of the human forebrain to demonstrate how HCMV infection induces syncytia, drives neurite retraction, and remodels microtubule networks to promote viral production and release. We establish that HCMV downregulates microtubule associated proteins at 14 days postinfection while simultaneously sparing other cytoskeletal elements, and this includes HCMV-driven alterations to microtubule stability. Further, we pharmacologically modulate microtubule dynamics using paclitaxel (stabilize) and colchicine (destabilize) to examine the effects on neurite structure, syncytial morphology, assembly compartment formation, and viral release. With paclitaxel, we found improvement of neurite outgrowth with a corresponding disruption to HCMV-induced syncytia formation and Golgi network disruptions but with limited impact on viral titers. Together, these data suggest that HCMV infection-induced disruption of microtubules in human cortical neurons can be partially mitigated with microtubule stabilization, suggesting a potential avenue for future neuroprotective therapeutic exploration.

**IMPORTANCE:** Infection by human cytomegalovirus (HCMV) continues to cause significant damage to human health. In the absence of a vaccine, vertical transmission from mother to fetus can result in profound neurological damage impacting quality of life. These studies focus on understanding the impact of HCMV infection on forebrain cortical neurons derived from iPSCs. We show that infection results in loss of neurite extension accompanied by cell-to-cell fusion. These pathogenic changes involve HCMV infection-mediated disruption to the microtubule network. Upon addition of the microtubule stabilization agent paclitaxel, the structural damage was limited, but infection still progressed to produce infectious particles. This work is part of our continued efforts to define putative strategies to limit HCMV-induced neurological damage.

## INTRODUCTION

Human cytomegalovirus (HCMV) is a pervasive pathogen that is estimated to infect between 40-90% of adults worldwide. (1, 2) As a human betaherpesvirus, HCMV infection is lifelong, occurring in waves of dormancy (latency) and reactivation. Further, transfer of infection can occur either through post-natal exposure to infected bodily fluids (horizontal transfer) or vertically from parent to fetus *in utero*. HCMV infects a wide range of cell types, including fibroblasts, various subtypes of endothelial and epithelial cells, placental trophoblasts, and hematopoietic precursors. (3–7) Neural cells have also been established as a site of infection. Neural progenitor cells (NPCs) are central nervous system-specific stem cells and a key site of infection within the human brain. (8–10) Other cell types demonstrated to sustain infection include astrocytes (11), microglia (12), oligodendrocytes (13), and ependymal cells (14). The potential for neurons to become infected has, however, been debated within the field. (11, 15–17) Recently, both our group and others have demonstrated that terminally differentiated neurons are susceptible to infection, regardless of whether their progenitors were induced pluripotent stem cell (iPSC) or fetal stem cells. (18–21) Further, infection has been demonstrated to have functional impacts on neurons, including alterations to calcium signaling and reduced action potential generation. (18, 21) While these studies have been helpful in characterizing HCMV-induced functional changes in neurons, further evaluation is needed to understand how these cells change structurally.

The neuronal cytoskeleton is composed of three primary elements: actin microfilaments, neurofilaments, and microtubules. Together, these structures function to resist structural deformation, maintain intracellular organization, and allow the cell to interact with its surrounding environment. Actin microfilaments are composed of polar, filamentous actin (F-actin) strands that offer a quick means for neurite expansion and retraction in neurons. Typically, actin structures are found near the cell membrane and enable dynamic interaction with the surrounding environment via filopodia and lamellipodia formation. (22–24) Synaptic structures are also organized specifically by actin microfilaments. (23) Finally, actin filaments are also capable of inducing cell movement via the associated motor protein myosin. Neurofilaments (NFs) are the neuron-specific members of the intermediate filament protein family. In comparison to actin microfilaments (and microtubules), NFs are not polar, and, therefore, have no associated motor proteins. NFs primarily serve to maintain cell structure and, secondarily, facilitate neuronal function. At axonal projections, NFs arrange to increase radial diameter, improving conductivity of electrical depolarizations (action potentials). (25–28) Finally, microtubules (MTs) compose the third branch of the neuronal cytoskeleton and perhaps serve the most varied role in neuronal physiology.

Microtubules are hollow, polar, cylindrical structures that are composed of tubulin proteins. Tubulins are a protein superfamily with three primary variants: alpha, beta and gamma. α- and ß-tubulins form heterodimers that are the primary components of microtubules, with structure arising from repeating, organized assembly of the dimerized proteins. (29) αß dimers bind guanosine diphosphate (GDP) or triphosphate (GTP) functional groups that dictate MT assembly. (30, 31) MTs are further strengthened through the binding of microtubule-associated proteins (MAPs) such as tau, MAP2, and MAP4. MAPs bind MTs longitudinally, acting both to crosslink individual heterodimers and connect cytoskeletal elements (MT-to-MT or MT-to-actin). (32, 33) At the negative end of the MT, αß dimers are stabilized using a combination of γ-tubulin and negative-end binding proteins. (34) γ-tubulin also plays a key role in nucleation for MTs at the centrosome (or, to a lesser degree, at the Golgi Apparatus), coordinating with other gamma-tubulin complex components (GCPs). (35, 36) MT nucleation events occur at a microtubule organizing center (MTOC). These structures act as a hub for MT extension throughout the whole cell and serve a key role in coordinating MT function.

Microtubules act as intracellular “highways”, facilitating the movement of cellular components to their proper destinations. This process is facilitated by MT-associated motor proteins such as dynein and kinesin to move cargo. This cargo can wildly range in size, including objects as small as viral particles and as large as entire organelles. (37) In neurons, MTs are indispensable for trafficking of neurotransmitter-containing synaptic vesicles (38) and facilitating energy distribution via trafficking of mitochondria (39). Additionally, MTs are key to maintaining neuronal morphology, whether semi-statically at the soma and axon or dynamically within dendrites. (40) Outside of these roles, MTs also have a putative role in intracellular signaling cascades. (41)

While HCMV’s effects on the neuronal cytoskeleton have not yet been evaluated, some insight can be derived from the virus’ effects on fibroblasts and other non-neural cells. Upon entry, viral particles utilize microtubules and dynein to traffic capsids from the cell membrane to the nucleus. (42–44) HCMV also reorganizes protein processing/exocytosis machinery (Golgi, trans-Golgi network, endosomes) into an infection specific organelle termed the viral assembly compartment (AC). (45–48) This structure functions as an MTOC within infected cells, effectively displacing the role of centrosomes within healthy cells. (49) MTs extending outward from the AC enable trafficking of both viral capsids/tegument proteins to the AC (using dynein) and movement of completed virions from the AC to the plasma membrane for exocytosis (via kinesin). (43, 45–47) Others have documented changes in microtubule-associated proteins upon HCMV infection, including increased expression of both γ-tubulin and end-binding protein 1 (EB1) and higher levels of EB3 activity at MT plus ends. (49) Interestingly, HCMV infection in fibroblasts demonstrated a decrease in MT structures by immunofluorescence as early as 12 hours post infection (hpi). (50)

Here we examined how HCMV alters the neuronal cytoskeleton in the context of iPSC-derived forebrain neurons. We observed significant structural alterations in neuronal morphology occurring between 2- and 14-days post infection (dpi) including shortened neuronal processes, reorganization of tubulin within syncytia, and areas of lower neurite density. However, only microtubule-associated protein (MAP) transcripts and protein showed significantly reduced expression throughout infection as compared to modest to no changes in expression of other structural components. Therefore, we next asked what impact modulating microtubule stability had on neurite structure in HCMV infection. We treated neurons with the microtubule stabilizing agent paclitaxel or the destabilizing agent colchicine and found that paclitaxel altered syncytial structure, modestly increased neurite length in HCMV infected cultures relative to a DMSO control, changed AC morphology, and slightly reduced viral titers. In contrast, colchicine had no discernable effect on syncytia shape, neurite structure, AC formation, or viral titers. Taken together, these data demonstrate that HCMV alters tubulin structures to generate abnormal morphology in infected neurons. Further, treatment of HCMV infected neurons with paclitaxel restores some degree of neuronal structure and may decrease overall viral output in infected neurons.

## RESULTS

### HCMV infection alters localization of tubulin and expression of microtubule-associated genes

We used four independent and unrelated iPSC lines to assess the impact of HCMV infection on human neuron structure. To generate forebrain neurons, we differentiated iPSCs into a neural progenitor cell (NPC) stage and subsequently patterned NPCs toward a forebrain neuron lineage **(Fig. 1A)**. Consistent with our previous work (18), neurons were infected with HCMV clinical strain TB40/e-eGFP at a multiplicity of infection (MOI) of 3 infectious units per cell at day 51 of differentiation **(Fig. 1B)**. Using a genetically encoded eGFP reporter, we determined sites indicative of viral infection **(Fig. 1C)**.

**Figure 1.**
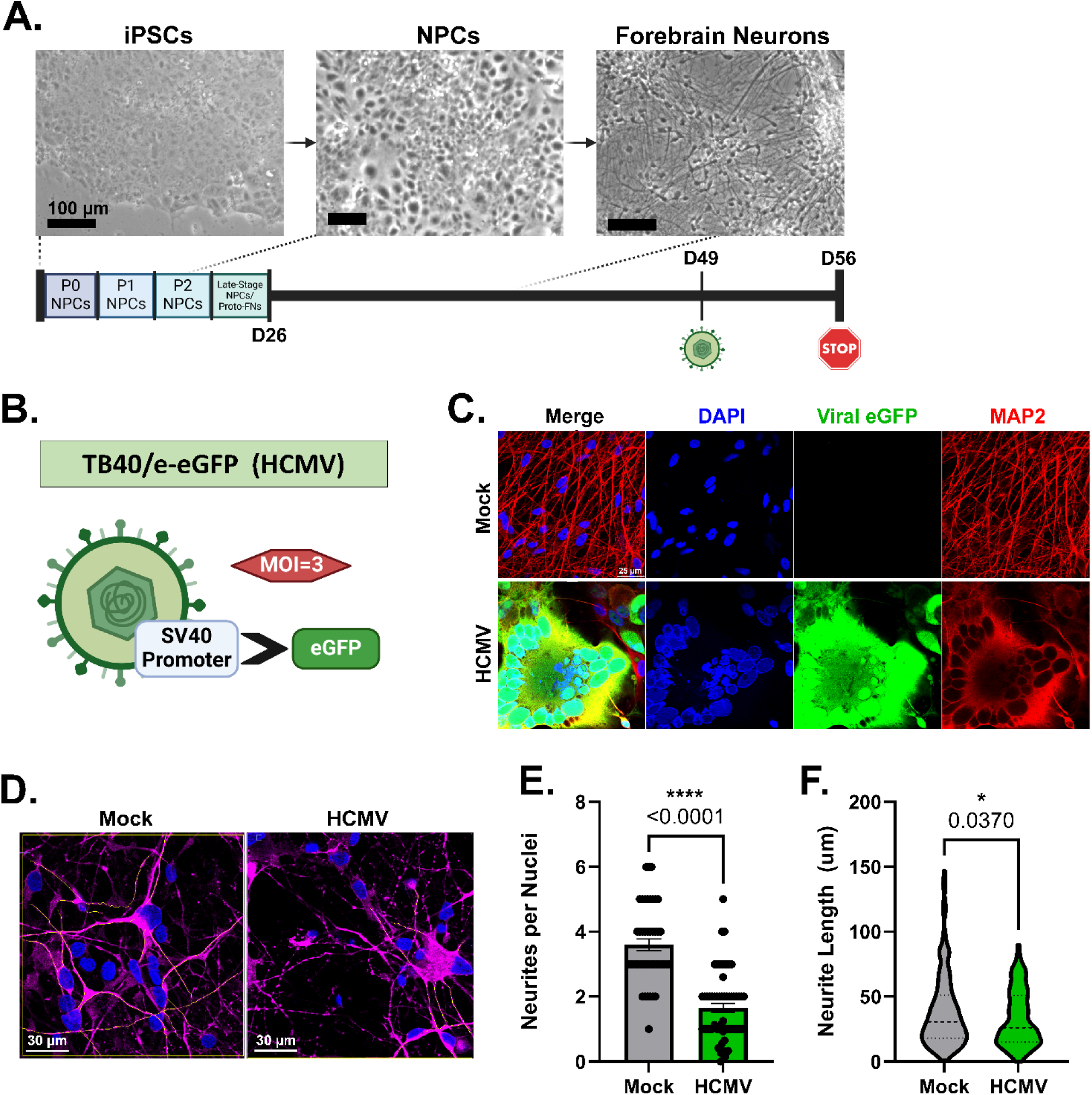
iPSC-derived forebrain neurons infected with HCMV demonstrate altered neurite morphology. (A) Schematic demonstrating derivation of forebrain neurons from iPSCs. (B) HCMV clinical variant TB40/e-eGFP encodes an eGFP fluorescent indicator controlled by an SV40 promoter. Neurons infected in these experiments were do so at an MOI of 3. (C) Representative image of mock-treated and HCMV-infected forebrains demonstrating typical infection-induced syncytial morphology (scale bar = 25 µm). (D) Representative images of manual neurite traces (yellow) using MAP2 staining in mock- and HCMV-treated forebrain neurons (scale bar = 30 µm). (E-F) Neurite metrics analyzed using manual tracing, including neurite per cell nucleus and neurite length measures. All data are presented as mean ± SEM. Unpaired Student’s t-tests were used to analyze statistical differences in E-F. *P < 0.05, **P < 0.01, and ***P < 0.001.

Initial observation of infected cultures revealed significant alteration to neuronal structure, including formation of syncytia and a visual change to neurites **(Fig. 1C)**. Using neurite tracing analysis in combination with MAP2 staining, we found that both neurite quantity per nuclei and neurite length were significantly decreased when neurons were infected with HCMV compared to mock conditions **(Fig. 1D-F)**. To further investigate this phenomenon, we sought to evaluate the effects of HCMV on various elements of the neuronal cytoskeleton, including actin filaments, neurofilaments and microtubules **(Fig. 2A)**. Assessment of gene expression for these structural components was first conducted via qRT-PCR. The effects of HCMV on ß-actin transcripts showed no change at 7 dpi but a significant decrease at 14 dpi **(Fig. 2B-C)**. Neurofilament subunits showed no significant HCMV-induced change in both light (NEF-L) and medium (NEF-M) chain mRNAs at both 7 and 14 dpi, but there was an HCMV-induced upregulation of heavy chain (NEF-H) transcripts at both timepoints **(Fig. 2D-E)**. Finally, mRNAs for neuron-specific beta-III tubulin (TUBB3) were found to be significantly downregulated at both 7 and 14 dpi **(Fig. 2F-G)**. These findings indicated potential downregulation of both actin and tubulin expression.

**Figure 2.**
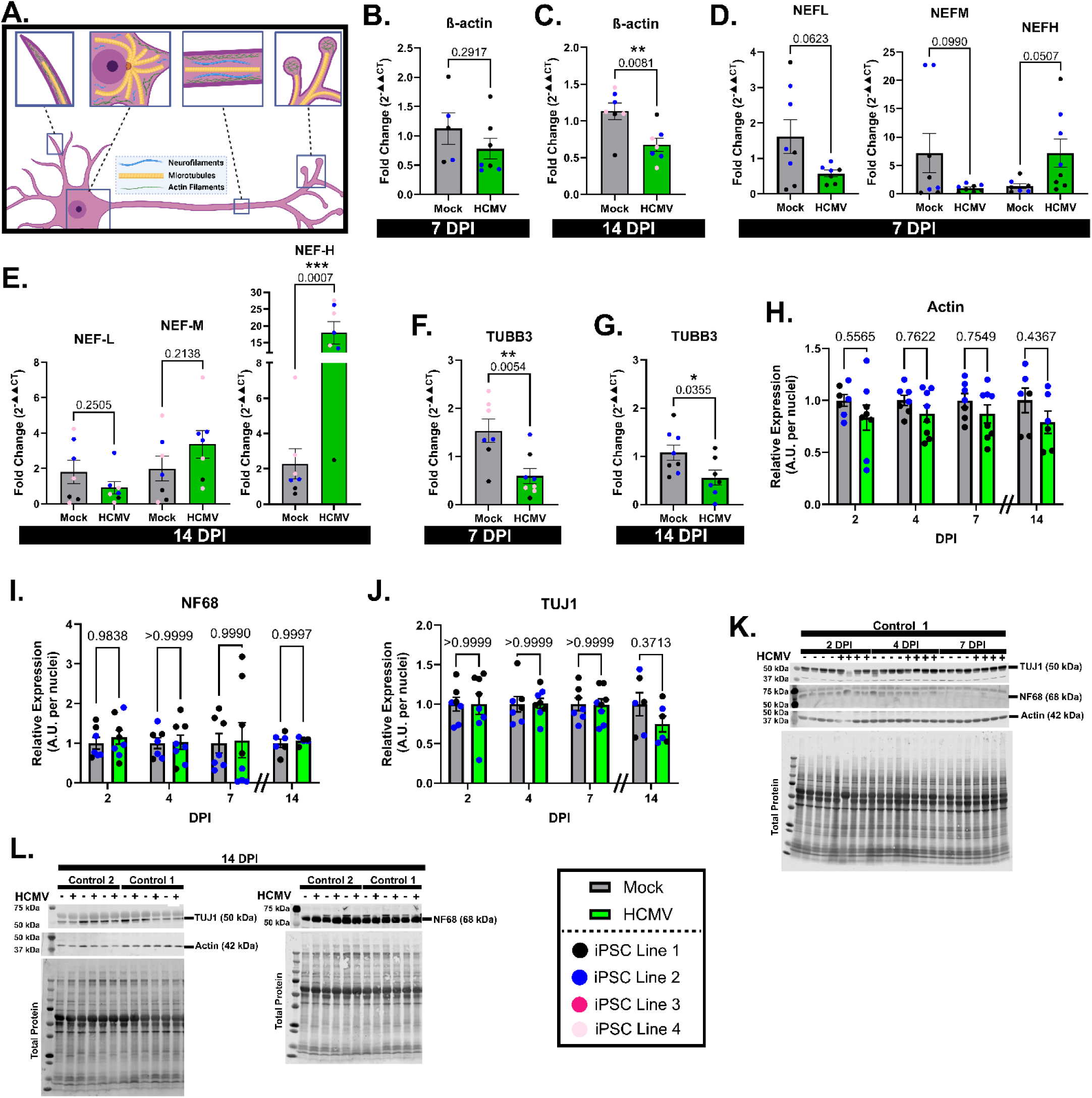
HCMV Effects on cytoskeletal transcripts and proteins. (A) Schematic showing the various components of the neuronal cytoskeleton. (B-C) HCMV demonstrates significant downregulation of actin mRNA at 14-days post infection (dpi), but not at 7 dpi. (D-E) Infection of forebrain neurons does not consistently alter neurofilament transcripts at either 7 or 14 dpi. (F-G) TUBB3 (neuron-specific ßIII tubulin) gene expression is significantly downregulated upon infection at 7 and 14 dpi. (H-L) Western blot quantification and images demonstrating that HCMV does not induce significant changes in actin, neurofilament 68 (NEFL) or tubulin protein expression at 2, 4, 7, and 14 dpi. All data are presented as mean ± SEM. Student’s t-tests were used to analyze statistical differences in 2B-2G. Ordinary 2-way ANOVA with Šídák multiple comparisons test (single pooled variance) was utilized for 2H-J. *P < 0.05, **P < 0.01, and ***P < 0.001.

We next aimed to examine the temporal changes in cytoskeleton elements at a protein level. We noted a consistent trend toward downregulation across all timepoints for ß-actin expression **(Fig. 2H, 2K-L)**, which mirrored a similar trend to the mRNA levels. Neurofilament 68 (NF68, NEF-L gene product) was selected to represent the effects of HCMV on neurofilament expression. There was no significant effect of HCMV on NF68 protein expression across all tested timepoints (2, 4, 7 and 14 dpi) **(Fig. 2I, 2K-L)**. Finally, neuron-specific ß-III tubulin (antibody clone TUJ1) was examined to assess HCMV’s effects on microtubules (MT). No significant change in TUJ1 expression was noted at any timepoint, though a modest decrease in tubulin was present at 14 dpi **(Fig. 2I, 2J-K)**. As we see HCMV-induced structural deformations of the neuronal cytoskeleton at 7 dpi **(Fig. 1C)**, these findings indicated that HCMV modulation of the cytoskeleton does not occur primarily through altered expression of structural proteins. Therefore, we next sought to evaluate proteins that modulate cytoskeletal stability.

We next used qRT-PCR to assess two key MAP genes: MAP2 and MAPT. MAPT transcripts were found to be downregulated at both 7 (∼48% decrease) and 14 dpi (∼62% decrease) **(Fig. 3A-B)**. Likewise, MAP2 transcripts were decreased at both timepoints by ∼72% (7 dpi) and ∼53% (14 dpi) **(Fig. 3C-D)**. Assessment of total tau protein (MAPT-encoded isoforms) by western blot found significant downregulation at both 7 (∼24%) and 14 dpi (∼50%) **(Fig. 3E-H)**. MAP2 protein expression was decreased by ∼31% at 14 dpi **(Fig. 3I-J)**. Together, these data suggest that neuronal microtubule stability may be impacted by HCMV infection.

**Figure 3.**
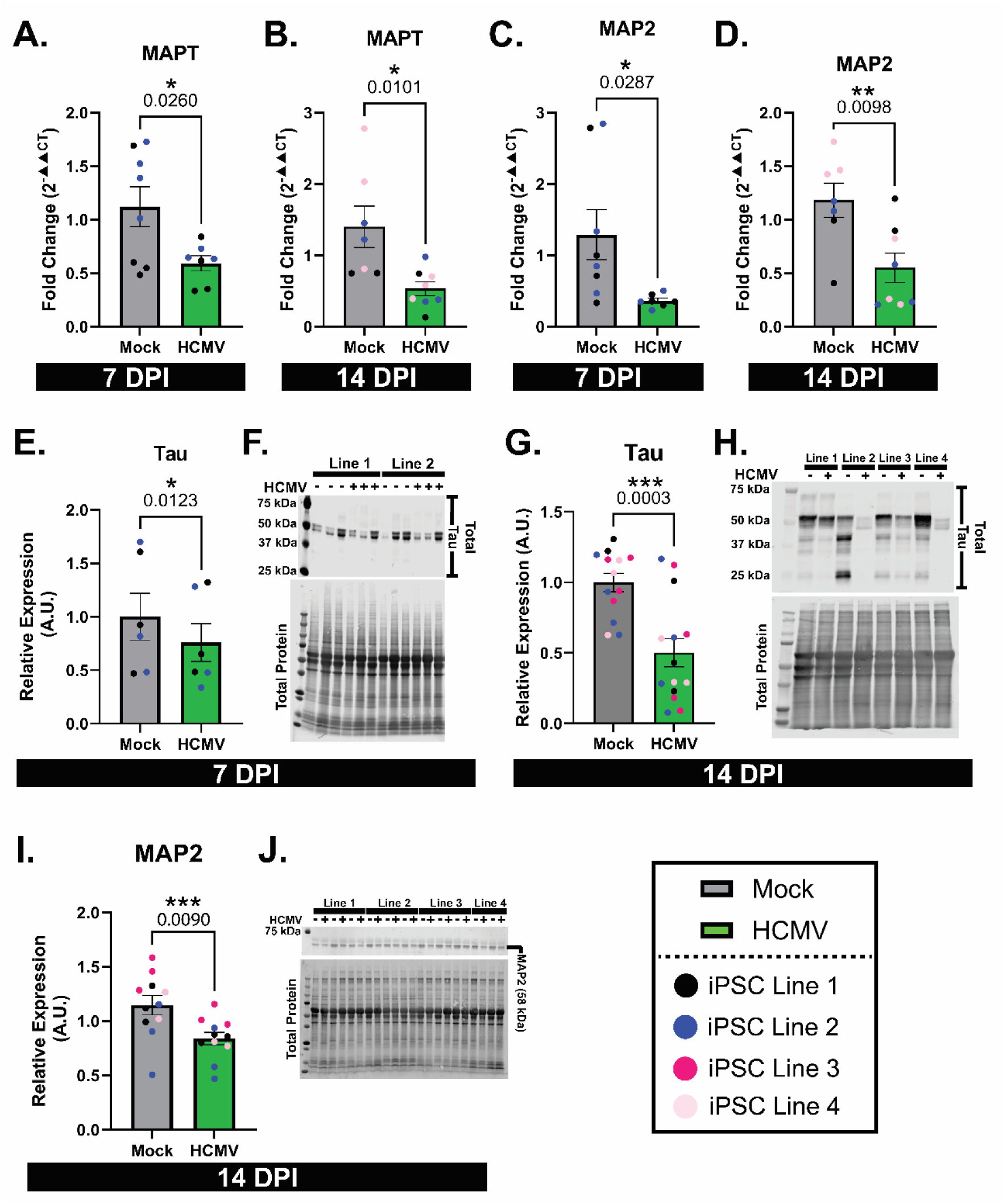
Infection with HCMV alters microtubule associated protein (MAP) expression. (A-B) MAPT transcripts are downregulated by HCMV at both 7 and 14 dpi timepoints. (C-D) HCMV derives significant decreases in MAP2 mRNA at 7 and 14 dpi. (E-H) Western blot quantification and images demonstrated HCMV-dependent decreases in tau (MAPT gene product) expression upon HCMV infection at both tested timepoints. (I-J) Infection induces MAP2 protein decreases at 14 dpi. All data are presented as mean ± SEM. Student’s t-tests (paired, where applicable) were used to analyze statistical differences. *P < 0.05, **P < 0.01, and ***P < 0.001.

Although there were no significant changes noted in tubulin expression through the first 7 days of infection, neuron structure was altered. We examined neuronal infection at 2, 4 and 7 dpi using immunofluorescence and observed changes in tubulin and MAP distribution within infected cells. First, a subclass of neurites associated with eGFP labeled cell bodies appeared thickened relative to their mock-treated counterparts **(Fig. 4A)**. This structural change was noticed as early as 4 dpi and continued to be observed within 7 dpi cultures **(Fig 4A, white arrowheads)**. Further, at 7 dpi, TUJ1 staining within syncytia revealed non-filamentous, punctate signal within the syncytial core **(Fig 4A, blue arrowheads)**. Finally, we noted areas devoid of neurite processes within infected cultures, especially distal to syncytia **(Fig 4A, orange arrowheads)**. These data suggested that HCMV induces a structural phenotype by utilizing the existing tubulin present within a cell. Therefore, we hypothesized that HCMV alters MT stability rather than affecting overall MT abundance.

**Figure 4.**
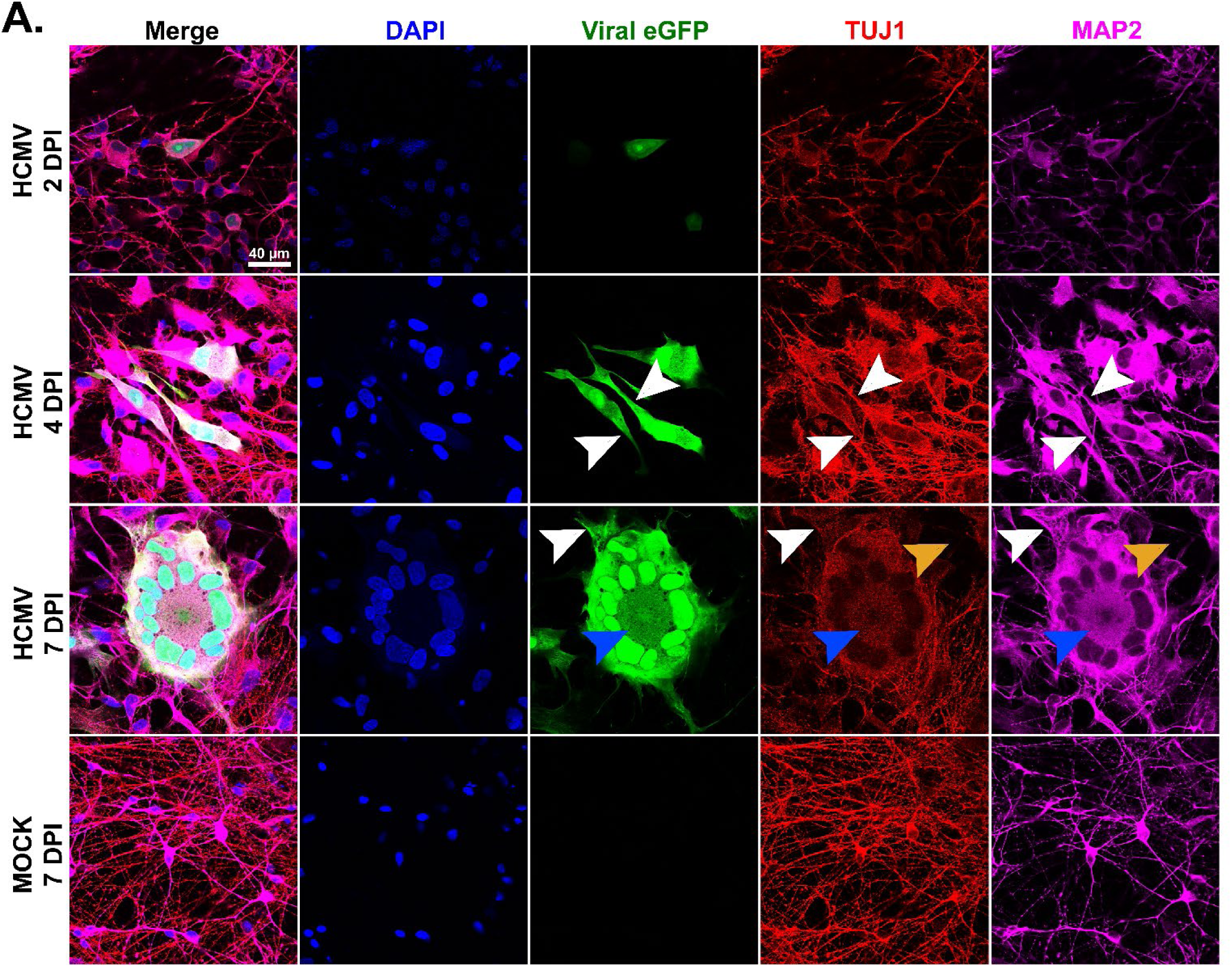
Time course evaluation of HCMV over the first 7 days of infection. (A) Forebrain neurons infected with HCMV begin to show eGFP-positive plaques as early as 2 dpi. Further, neurite changes can be observed as early as 4 dpi relative to mock cultures. These findings include thickened processes (denoted by white arrows). At 7 dpi, tubulin and MAP2 staining appeared punctate in the core of syncytia (blue arrows), rather than filamentous as examined elsewhere. Further, regions lacking neurites were more common in the areas adjacent to the syncytia (orange arrows). Scale bars = 40 µm

### Stabilization of microtubules with paclitaxel alters syncytial morphology

We next utilized two compounds known to impact the formation and stability of MTs to determine the role of microtubule stability in forebrain neuron infection: paclitaxel, an MT stabilizing agent, and colchicine, an MT destabilizing compound. As microtubules are required for HCMV entry and pathogenesis, treatment was delayed until 24 hours post infection, and subthreshold amounts of each compound were used to avoid the excessive cell death observed at higher dosages. (51–54) Neurons were treated with paclitaxel (Pac), colchicine (Col), or DMSO (vehicle) for 6 days before collection of cultures at 7 dpi **(Fig. 5A)**. Initial assessment of TUJ1 was conducted at 7 dpi to determine if drug application altered expression. As expected, no significant differences were detected between the various treatment groups **(Fig. 5B-C)**. Following this, we investigated tubulin structure via immunofluorescence. Within mock treated cultures, we detected no significant differences in neurite quantity or length, though Pac treated cultures were non-significantly decreased relative to DMSO controls (p=0.1350) **(Fig. 6A, 6C, 6E)**. Immunofluorescence demonstrated that syncytia formed in the untreated and DMSO-treated infected cultures and were similar in shape **(Fig. 6A-B)**. Further, measurements of neurite number and neurite length were not significantly different between these groups **(Fig. 6D, 6F)**. Interestingly, paclitaxel-treated neurons formed syncytia of a novel shape (linear rather than circular) **(Fig. 6A-B)**. Further, infected neurons treated with paclitaxel trended toward having longer neurite lengths relative to DMSO **(Fig. 6D),** but overall neurite counts were not affected **(Fig. 5F)**. Colchicine had no discernable effect on neurite count **(Fig. 6F)**, syncytia shape **(Fig. 6A-B)**, or neurite quantity and length **(Fig. 6D, 6F)**. These data indicated that stabilization of microtubules can disrupt typical syncytia organization induced during HCMV infection. Considering this, we hypothesized that Pac could also disrupt the formation of the viral assembly compartment.

**Figure 5.**
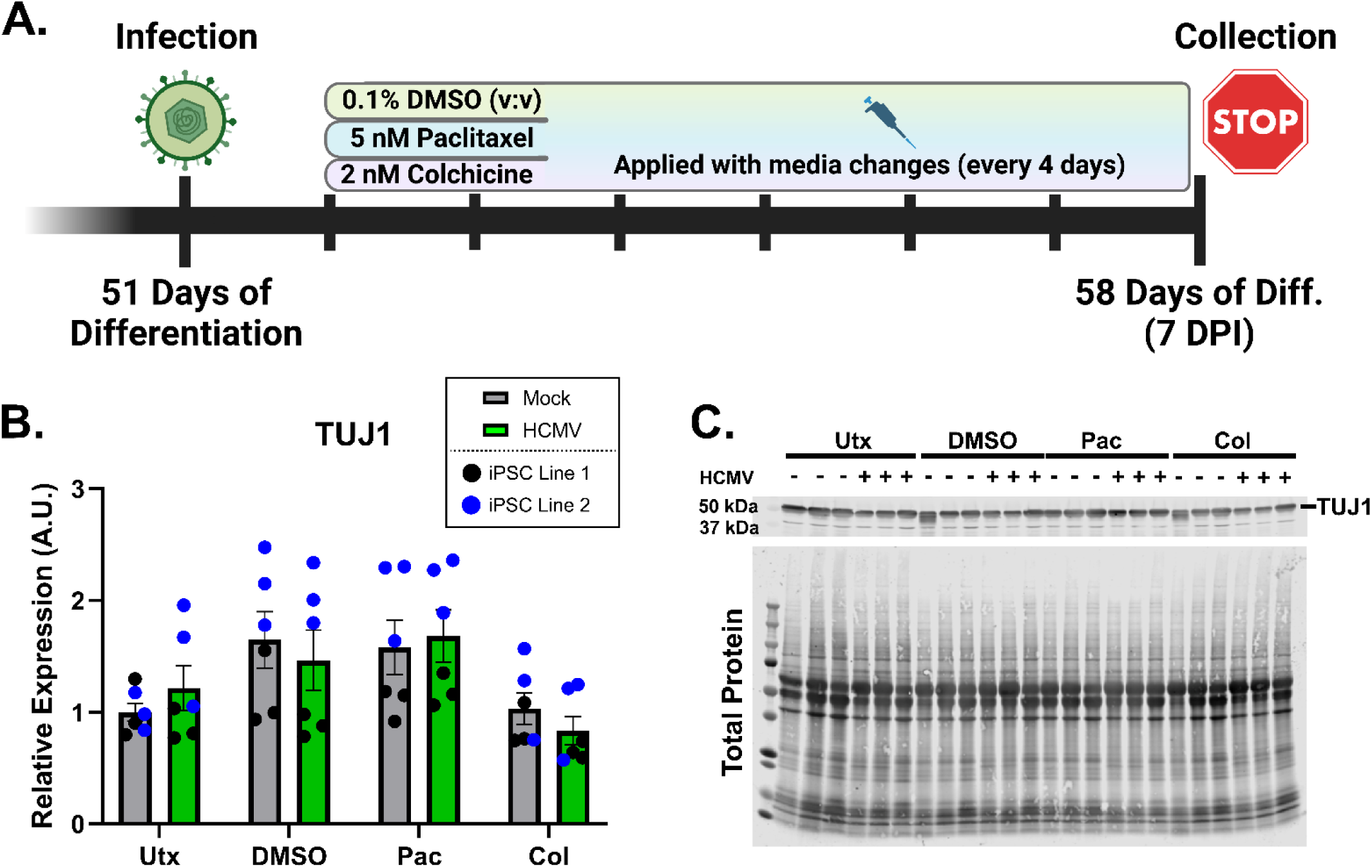
Paclitaxel and colchicine application to alter microtubule stability in infected forebrain neurons. (A) Diagram of the dosing schedule for paclitaxel, colchicine, and DMSO vehicle for application to forebrain neurons. (B-C) Western blot analysis and image for neuron-specific beta III tubulin (TUJ1) expression. Application of DMSO and microtubule-affecting drugs does not appear to impact expression of tubulin. All data are presented as mean ± SEM. An ordinary 2-way ANOVA with Tukey’s post-hoc analysis was used to analyze statistical differences in 5C. *P < 0.05, **P < 0.01, and ***P < 0.001.

**Figure 6.**
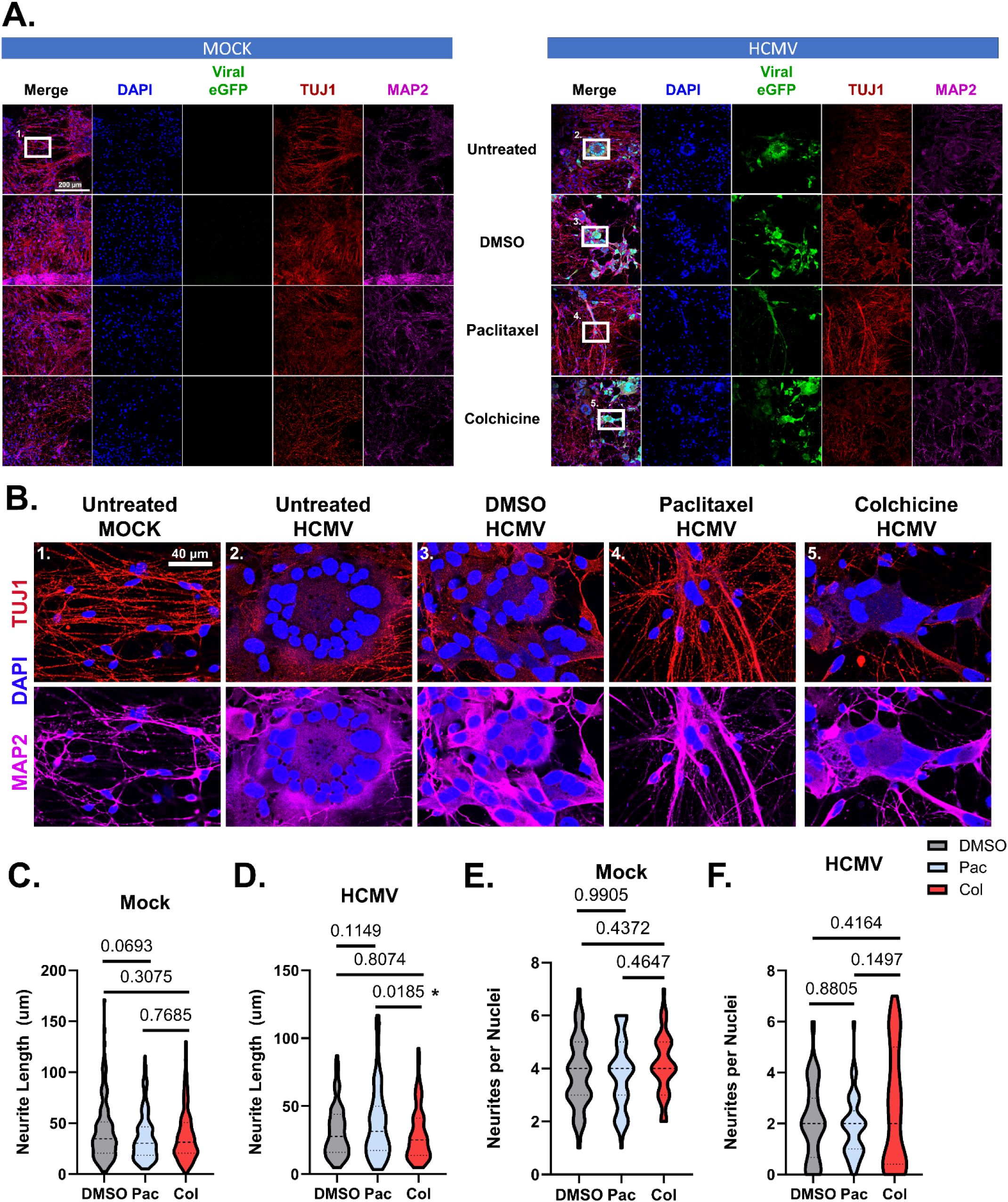
The effects of paclitaxel and colchicine on neurite structure in HCMV-infected forebrain neurons. (A) Large-format images (500 µm x 500µm) detailing the effects of DMSO vehicle, paclitaxel, and colchicine on neurite structure. White boxes are ROIs that are magnified in 3.5B. Scale bar = 200 µm (B) Zoomed ROIs from 3.5A demonstrating tubulin and MAP2-labelled neurites upon treatment with Pac, Col and DMSO. Paclitaxel demonstrates a partial restoration of neurites by IF. Scale bar = 40 µm. (C-D) Quantification of neurite length (using MAP2 staining) in both mock (C) and HCMV (D) conditions. (E-F) Evaluation of the effects of Pac, Col and DMSO on number of neurites per nucleus in both mock (E) and infected (F) conditions. All data are presented as mean ± SEM. Ordinary 1-way ANOVA and Tukey’s post-hoc analysis used to analyze statistical differences in C-F. *P < 0.05, **P < 0.01, and ***P < 0.001.

### Viral assembly compartment morphology is changed in paclitaxel-treated neurons

HCMV infection induced formation of an AC within neurons, as indicated by trans-Golgi network protein 46 (TGN46) staining **(Fig. 7A)**. TGN46 and γ-tubulin were each assessed to examine any drug-related alterations to expression, and neither was altered upon addition of paclitaxel and colchicine **(Fig. 7B-D)**. Having established no significant differences in AC and MT-nucleating proteins, AC structure was visualized using immunofluorescence **(Fig. 8A-B)**. In mock-treated cells, neurons exposed to DMSO, Pac, and Col looked identical to untreated cells and were excluded from **Figure 8** as a result. In infected cultures, DMSO treatment was not found to substantially alter AC formation relative to untreated cells **(Fig. 8A-B)**. However, Pac-treated syncytia exhibited long, smear-like ACs running the length of the structure, next to the linearly arranged nuclei **(Fig. 8A-B)**. Col-treated infected neurons, while still demonstrating round ACs, appeared smaller than those formed by other treatment conditions **(Fig. 8A-B)**. Together, these data suggest that paclitaxel treatment is sufficient to disrupt HCMV-induced cell morphology and syncytia structure, which led us to consider effects of Pac treatment on viral production.

**Figure 7.**
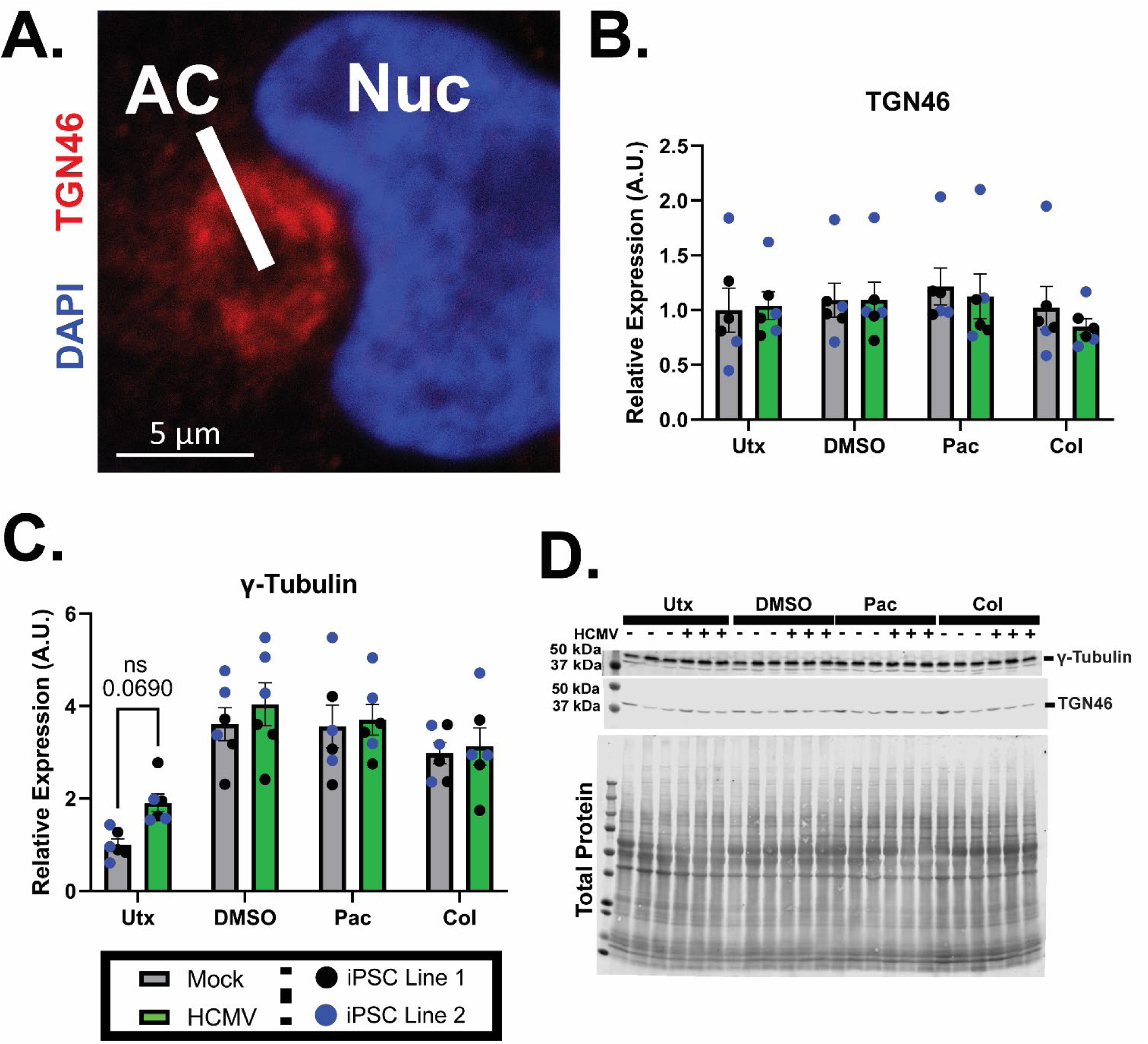
Drug treatments do not substantially alter the materials used to nucleate microtubules in forebrain neurons. (A) Confocal image of the viral assembly compartment (AC) next to the nucleus of an infected neuron. Scale bar = 5 µm. (B, D) Western blot analysis of TGN46 (a marker of the HCMV AC) reveals that protein concentration is not affected by either HCMV infection or treatment with Pac, Col or DMSO. (C-D) Evaluation of γ-tubulin (a nucleating unit for microtubules) demonstrates similar results, though untreated cells do display a near-significant increase upon infection. All data are presented as mean ± SEM. Ordinary 2-way ANOVA with Tukey’s post-hoc analysis was used to analyze statistical differences in B-C. *P < 0.05, **P < 0.01, and ***P < 0.001.

**Figure 8.**
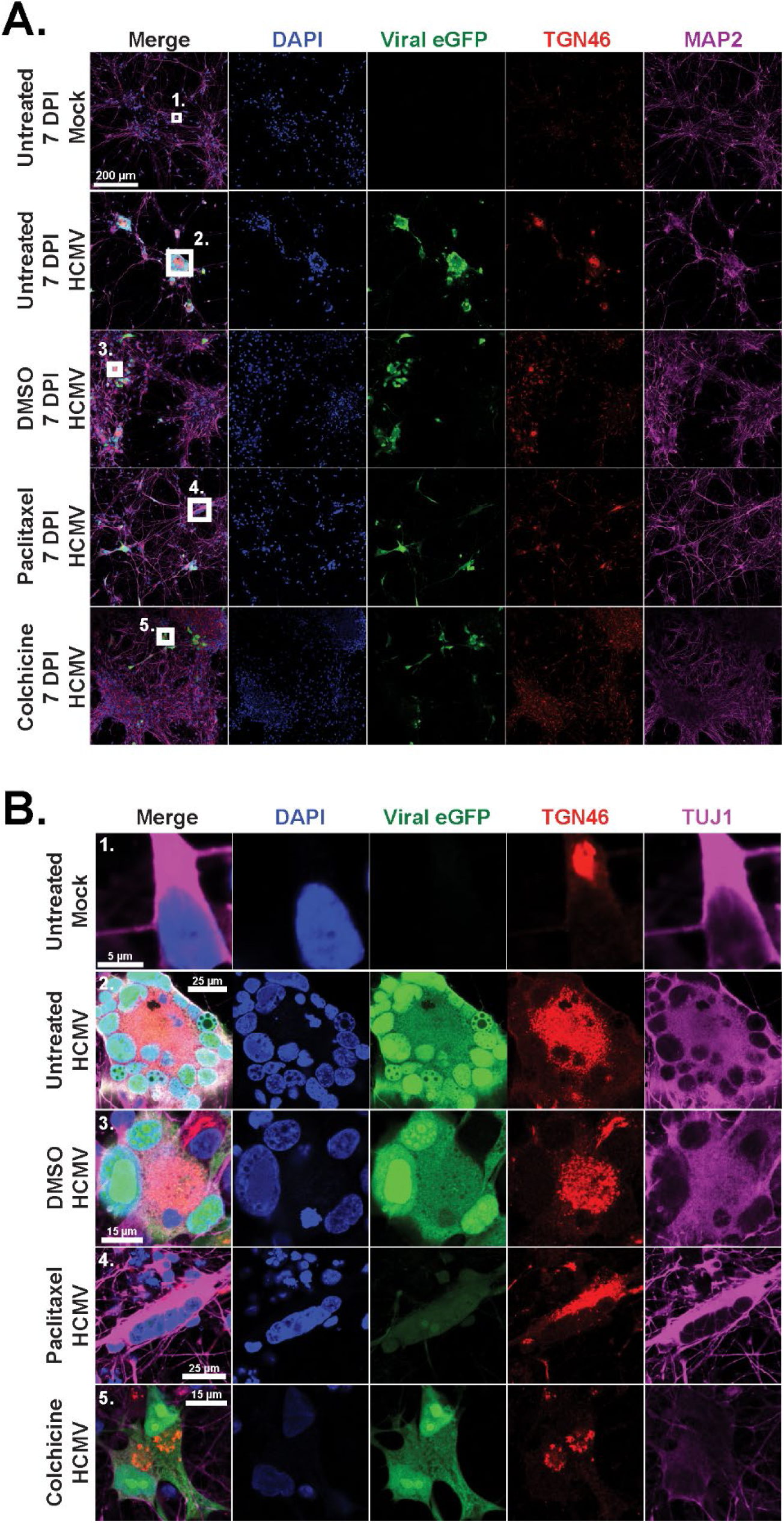
Effects of paclitaxel and colchicine on assembly compartment structure. (A) Large-area images (500 µm x 500 µm) of 7 dpi mock neurons or 7 dpi infected neuronal cultures treated with DMSO, paclitaxel, or colchicine. White boxes indicate inset images. Scale bar = 100 µm. (B) Inset images from 3.7A highlighting individual cells or syncytia to visualize their trans-Golgi (mock) or AC (infected). Paclitaxel-treated cells demonstrate irregular AC formations relative to untreated infected neurons. Scale bars are variable and noted in each image.

### Markers of viral production are not significantly altered by modulating microtubule stability

As AC complexes were affected by Pac treatment, we next sought to examine the subsequent impact of MT stabilization modulation on viral output from infected neurons. Live imaging was conducted for ∼7 dpi, with frequent monitoring of eGFP expression within the culture. These traces are depicted in **Figure 9A** and did not demonstrate any significant deviations by Pac or Col treatment relative to DMSO-treated and untreated controls. These findings were confirmed upon evaluation of total eGFP expression in 7 dpi cell lysates **(Fig. 9B-C)**. Cell lysates were also probed for relative concentrations of HCMV immediate early protein 1 (IE1) and late protein pp28. No significant differences were detected between any of the treatment groups for these targets **(Fig. 9D-F)**. Finally, viral titers were analyzed using the conditioned medium from HCMV-infected neurons. There were no significant differences between treatment groups, but Pac treated cells did trend toward reduced viral titers **(Fig. 9G)**. Taken together, these data suggest that HCMV-induced structural changes can be partially abated by paclitaxel treatment with a correspondingly subtle reduction in viral production.

**Figure 9.**
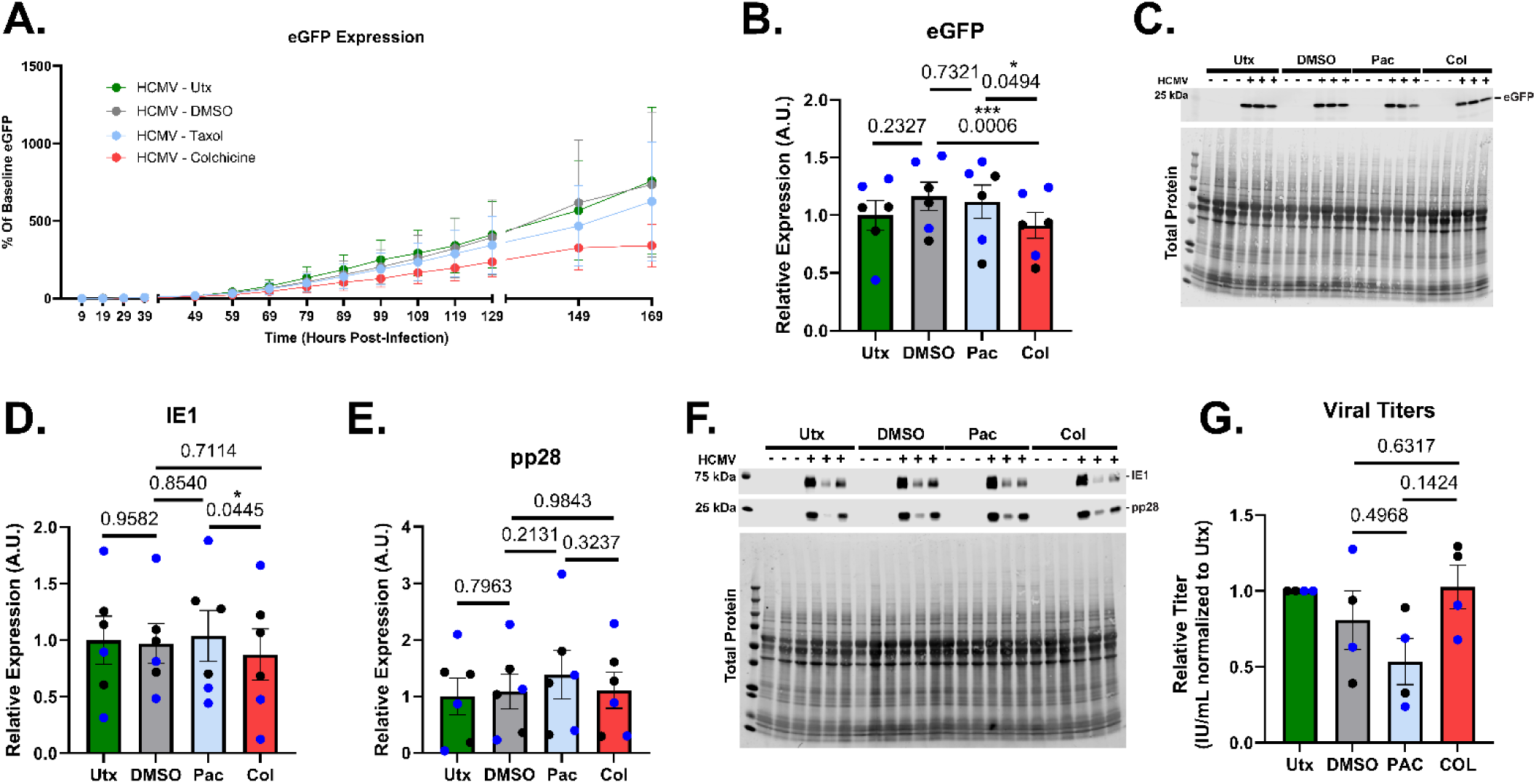
The effects of paclitaxel and colchicine treatments on viral output in HCMV-infected forebrain neurons. (A) Viral eGFP expression data was collected over the first 169 hpi for neurons treated with nothing, DMSO, paclitaxel, and colchicine. No significant differences were noted in the amount of eGFP between conditions. (B-C) Likewise, western blot analysis of eGFP was found to be consistent, with no discernable differences between treatment groups. *Note*: this blot was stripped of antibody and reused, so the total protein stain is repeated from Figure 5C. (D-F) Immediate early gene 1 protein (IE1) and pp28 are markers of viral infection, but neither are significantly altered with treatment. (G) Finally, viral titers were assessed for each condition, revealing no significant changes between DMSO, paclitaxel and colchicine, though a trend toward reduced viral production was noted in Pac-treated cells. All data are presented as mean ± SEM. Ordinary 1-way ANOVA with Geisser-Greenhouse correction and Tukey’s post-hoc analysis was used to analyze statistical differences in A, B, D, E, G. *P < 0.05, **P < 0.01, and ***P < 0.001.

## DISCUSSION

Microtubules have been known to be associated with HCMV infection for decades. Their relevance in the virion entry, capsid trafficking, viral assembly, and, ultimately, egress processes have been thoroughly documented in various cell models (49, 55–59), contributing significantly to our understanding of HCMV as a pathogen. However, the recent groundswell of interest regarding HCMV’s effects in nervous tissue (19, 21, 60–66) has resulted in neurovirologist relying almost exclusively on reports conducted in fibroblasts and epithelial cells to draw conclusions about neural cells. Due to substantially different structures and functions, the application of findings from fibroblasts onto neurons and NPCs is likely to be problematic. Fortunately, since the discovery of iPSCs, access to human-derived neural cells has increased exponentially, allowing for easier evaluation of HCMV within a relevant model system. Here, we sought to better describe neuron-specific structural phenomena relating to HCMV infection. Specifically, we analyzed HCMV’s effects on expression of neuronal cytoskeletal components and demonstrated that stabilization of microtubules can mitigate some of the morphological aberrancies induced by infection.

HCMV induced significant alterations to neuronal morphology within the first 7 days of infection **(Fig. 1C, 4A)**. Neurites (axons and dendrites) have been described by both our group and others as a focal site of these changes, with projections becoming dissociated from their underlying matrix (21) and ultimately degenerating or retracting into the neuronal soma (18, 21) **(Fig. 1D-F)**. The other significant morphological change that occurs in HCMV infected neurons is the formation of multinucleated syncytia **(Fig. 1C)**. (18) While the latter’s mechanism is likely conserved from other cell types – viral proteins on the plasma membrane surface driving host cell fusion – little is understood about the way in which retraction of cellular projections occurs. Our assessment of cytoskeletal elements indicated that neurite loss is not due to altered gene expression, as neurofilaments, actin, and tubulin are not differentially expressed at 7 dpi **(Fig. 2H-L)**. Although, interestingly, transcripts for actin and tubulin appear to be downregulated at 7 dpi, this does not translate to the protein level **(Fig. 2B-L).** It is possible this is due to HCMV-induced altered host cell protein degradation (67), but more work is needed to address this. We also noted significant decreases in the expression of MAP2 at 14 dpi (∼25%) and tau at both 7 (24%) and 14 dpi (∼50%). Together, these findings suggest that the reorganization of the neuronal cytoskeleton might be enabled via altered MT stability rather than differential expression of subunits over the course of infection.

Utilizing both paclitaxel and colchicine, we sought to examine how pharmacological modulation of MT stability would impact HCMV-induced cytoskeletal rearrangement. Both compounds are FDA approved pharmaceuticals (68, 69) and have been proven efficacious for the stabilization (Pac) and destabilization (Col) of MTs in both in vivo and in vitro systems. (70–73) When deciding our concentrations for each drug, we wanted to ensure an appropriate balance between the compound’s intended effects and the chance for cytotoxicity. The upper limit for each compound was 10 nM due to documented cell death above that range. (51, 54) However, as we intended to incubate neurons with each for a minimum of 6 days, we decided upon concentrations well below those utilized in short term studies **(Fig. 5A)**. It remains possible that the modest effects we describe are due to this low dosage. For this reason, additional follow-up studies are necessary to better titer each drug for peak effectiveness. To ensure drug application was not aberrantly affecting gene expression, we conducted frequent “checkpoint” assessments on relevant proteins. From these we can conclude that Pac and Col do not significantly dysregulate TUJ1 **(Fig. 5B-C)**, TGN46 **(Fig. 7B, 7D)**, or γ-tubulin **(Fig. 7C-D)**. These findings imply that any changes induced by Pac and Col are due specifically to altered MT stability and not increased/decreased gene expression. However, Pac treatment did induce some restoration of neurite structure in treated cultures relative to the untreated infection group **(Fig. 6A-B)**. Quantification of neurite length in Pac-treated cells demonstrated a trend toward increased neurite length relative to infected neurons treated with DMSO **(Fig. 6D)**. Interestingly, while not significant, mock treated cultures demonstrated an opposing trend, with neurites appearing shorter relative to those in the mock-DMSO group **(Fig. 6C)**, which may be indicative of toxicity in otherwise healthy neurons. These data suggest that potential benefits of Pac treatment in the context of HCMV neuronal infection would need to be weighed with the potential negative effects to surrounding neural tissues.

In addition to the structural damage induced by HCMV, neurons undergo severe functional disruptions including reductions in calcium dynamics and elimination of action potential generation. (18, 60, 61) Additional research is needed to determine the impact of Pac treatment on calcium dynamics and electrophysiological activity in HCMV infected neurons to fully understand the potential benefits or consequences of MT stability in HCMV neuronal infection. However, considering the importance of neurite structure in neuronal signaling (74, 75), we would hypothesize that even a modest maintenance of structure could improve overall synaptic activity. In support of this, others have found that low-dose Pac positively impacts neuronal function in models of Alzheimer’s Disease (76), tauopathy (77), and spinal cord injury. (78)

At 7 dpi, Pac was found to alter syncytia formation **(Fig. 6A-B)**. Whereas typical neuronal syncytia are identifiable by their ringlet of nuclei and central cavity containing the AC (18), a subset of the structures in the Pac-treated neurons were found to have linear assembly of their nuclei **(Fig. 6A, 6B4)**. Initially, we hypothesized that this was resulting from Pac delaying syncytial formation. However, we never detected the presence of these linear syncytia in any of the other infected neuron conditions at 2 and 4 dpi. Similarly, we found Pac’s effects on the AC to be novel. Whereas a normal cell has a single trans-Golgi structure located proximally to the nucleus **(Fig. 8A-B, top rows)**, syncytia form a central AC derived from several cells’ worth of combined Golgi Apparatus, trans-Golgi, and early endosomes organized into concentric rings. (47–49) A subset of Pac-treated cells, however, displayed long, smear-like ACs along the length of their linearized syncytial structures **(Fig. 8A-B, fourth row)**. Interestingly, as noted in **Figure 7B-C**, no notable difference in quantity of TGN46 staining was found between Pac-treated and DMSO-treated/untreated controls. Together, these data indicate that Pac induced a specific syncytial morphology in treated HCMV-positive neurons. However, to our surprise, this altered syncytia structure did not translate into a significant disruption viral infection or viral production. Neither live imaging for eGFP expression **(Fig. 9A-C)** nor expression of the viral proteins IE1 and pp28 **(Fig. 9D-F)** showed a substantial reduction with Pac treatment. However, there was a trend toward reduced viral titers in the Pac treated samples **(Fig. 9G**). It is possible that with higher Pac concentration or a longer duration treatment paradigm there would be a more obvious decrease in viral titers, but that may come at the expense of overall neuron health. Nevertheless, even a modest reduction in viral titers and/or syncytia structure, in combination with improved neurite outgrowth, could have subsequent downstream beneficial effects on neuron health, survival, and function.

In this study, we describe structural changes in human iPSC-derived forebrain neurons infected with HCMV through the lens of cytoskeletal gene expression and microtubule stability. Paclitaxel is highlighted as a potential candidate for remedying the structural phenotype induced by infection in neurons. Further, the effects of modulating microtubule dynamics are seen on both syncytial and assembly compartment structure, which may have subsequent impacts on the overall infection. Together, this novel analysis of HCMV-induced cytoskeletal changes in human cortical neurons provides a foundation for further discoveries in this nascent field.

## METHODS

### Cell Culture and Differentiation

Induced pluripotent stem cell (iPSC) lines were derived from either reprogrammed skin fibroblasts or patient blood cells (Line 1 - Coriell GM03814 [Fibroblast line reprogrammed to iPSCs]; Line 2 – WiCell PENN022i-89-1 [purchased as iPSCs; derived from blood cells]; Line 3 – Coriell AG27607D [purchased as iPSCs; derived from fibroblasts]; Line 4 – Coriell AG25370 [purchased as iPSCs; derived from fibroblasts]). The utilized cell lines were chosen due to availability, consistency of differentiation, and diversity of patient background. Lines 1, 3, and 4 were derived from white females, with patient ages of 30, 69 and 81, respectively. Line 2 was obtained from a 28-year-old African American male. Neither we nor previous studies have evaluated HCMV infection status (latency) in these individuals prior to utilizing the above cell lines. Stem cell cultures were expanded under feeder-free culture conditions using Geltrex (Gibco) basement membrane matrix. iPSCs were plated onto 6-well plates and maintained with complete, daily replacements of Essential 8 medium (Thermo Fisher Scientific). Post-thaw, all iPSC cultures were passaged a minimum of 3 times before differentiation. All cultures were subject to quarterly mycoplasma testing to ensure freedom from contamination.

Patterning of iPSC colonies toward a neural progenitor (NPC) fate was completed utilizing dual-SMAD inhibition, as described by Chambers et al. (79) For this step, STEMdiff SMADi Neural Induction Kit (STEMCELL Technologies) and lab-prepared neural progenitor medium (Neurobasal, 10% v:v Knockout Serum Replacement [Gibco], 1% v:v 100x Non-essential Amino Acids [Gibco], 2% v:v 50X B27 Supplement [Gibco], 1% v:v 100x N2 Supplement [Gibco], 1% v:v 100x Antibiotic-Antimycotic [Gibco], 100 ng/mL Laminin [Sigma], 40 µM SB431542 [Biogems], 0.2 µM LDN193189 [Biogems] were used interchangeably. NPCs were cultured for a minimum of 3 passages in SMAD-inhibition medium (with daily changes) prior to differentiation to ensure complete and consistent generation of neural progenitors. For each passage, NPCs were dissociated using accutase (STEMCELL Technologies) to generate a single-cell suspension. Cells were then replated at a density of 2×10^5^ cells/cm^2^ onto Geltrex-coated 6-well plates. NPCs were passaged once every 6-7 days. Between 18-21 days of differentiation, NPCs were dissociated and plated at a density of 1.25×10^5^ cells/cm^2^ in preparation of patterning toward immature forebrain neurons.

Using the STEMdiff™ Forebrain Neuron Differentiation Kit (STEMCELL Technologies), NPCs were directed toward a forebrain neuron cell fate. Cells were maintained under daily media changes for 6-7 days prior to final accutase dissociation and plating onto either Poly-L-Lysine- and Laminin-glassware or Geltrex-coated plasticware at varying densities (6 well plate – 5.2×10^4^ cells/cm^2^, 24 well plate – 1.32×10^5^ cells/cm^2^, Coverslips – 3.1×10^5^ cells/cm^2^). Upon plating, immature forebrain neurons were cultured in forebrain neuron maturation medium (BrainPhys Neuronal Medium (STEMCELL Technologies), 2% v:v 50X B27 Supplement [Gibco], 1% v:v 100x N2 Supplement [Gibco], 1% v:v 100x Antibiotic-Antimycotic [Gibco], 100 ng/mL Laminin [Sigma Aldrich], 10 ng/mL Brain-Derived Neurotrophic Factor [PeproTech], 10 ng/mL Glial Cell Line-Derived Neurotrophic Factor [PeproTech], 1 µg/mL dibutyryl-cAMP [STEMCELL Technologies] until use.

### Viruses

Fluorescently labelled TB40/E (TB40/E-eGFP) was generously provided by Felicia Goodrum (University of Arizona, Tucson, AZ). This HCMV variant was generated via transfection of MRC-5 fibroblasts with both a bacterial artificial chromosome (BAC) containing the HCMV genome and a UL82-encoding plasmid, as previously described. (87–89) Utilizing the stocks of TB40/E-eGFP prepared in fibroblasts, ARPE-19 epithelial cells were infected and utilized to produce epithelial cell-derived TB40/E-eGFP. Viral titers were determined for epithelial-derived stocks using the method previously reported by our group (18, 80, 81) and are expressed in infectious units per milliliter (IU/mL).

All neuronal infections were conducted at a multiplicity of infection (MOI) of 3. Viral inoculum (or PBS for mock-treated) was applied to cell medium for 2 hours at 37°C, with constant agitation (rotary shaker plate). After exposure, cells were washed once with 1x Dulbecco’s PBS (dPBS, Gibco) to remove residual viral particles, and fresh media was added.

### Immunofluorescence and Live Imaging

Forebrain neuron cultures were plated onto 12mm Poly-L-Lysine (PLL, Sigma)/Laminin (Sigma)-coated coverslips at a density of 35,000 per coverslip (3.09×10^5^ cells/cm^2^). Cells were fixed in 4% paraformaldehyde for 15 minutes, washed once with dPBS, and stored in fresh dPBS at 4°C until use. Cell permeabilization was conducted by applying Triton X-100 solution (0.2% v:v, in PBS) to each coverslip and incubating for 10 minutes at room temperature (RT). After 1x PBS wash, cells were treated with blocking buffer (5% v:v normal goat/donkey serum [NGS/NDS; dictated by secondary] in PBS) for 1 hour (RT). Primary antibody solution (primary antibodies, 5% v:v serum [NDS/NGS], 0.1% v:v Triton X-100, in PBS) was then applied to coverslips overnight at 4°C. After 4x PBS washes, cells were treated with secondary antibody solution (secondary antibodies, 5% v:v serum, 0.1% v:v Triton X-100, in PBS) and allowed to incubate for an hour at RT. Secondary antibody solution was removed and coverslips were washed 4x with PBS to clear non-specific binding. Coverslips were mounted onto slides using Fluoromount-G Mounting Medium with DAPI (SouthernBiotech) and sealed using clear nail polish (Ted Pella Inc.) Coverslip imaging was conducted using a Zeiss LSM980 confocal microscope at various magnifications (20-63x). Images analysis was conducted using NIS Elements (Nikon), Zen Blue (Zeiss), and ImageJ.

The following antibodies and dilutions were used for immunofluorescence during this study: chicken anti-TUJ1 (1:250-300; GeneTex), chicken anti-MAP2 (1:250-500; Invitrogen), mouse anti-MAP2 (1:250-500, Invitrogen), rabbit anti-TGN46 (1:500, Invitrogen), goat anti-chicken IgY (H+L) AF647 (1:250, Invitrogen), goat anti-rabbit IgG (H+L) AF568 (1:500, Invitrogen), donkey anti-mouse IgG (H+L) (1:250, Invitrogen), and goat anti-chicken IgG (H+L) AF568 (1:500, Invitrogen).

Single-instance live cell imaging of virally encoded eGFP was collected using a Nikon TS100 inverted microscope. Additionally, timelapse live imaging was conducted using an IncuCyte S3 in-incubator system (Sartorius AG). Images were collected every 2 hours for 7 days, using settings to capture brightfield and green fluorescence images (ex. 440-480 nm; em. 504-544 nm). IncuCyte imaging was conducted using a 20x objective, and analysis was conducted using the IncuCyte 2022B revision 2 software package (video stitching, background subtraction, and eGFP quantification) and ImageJ (neurite retraction).

### Neurite Measurements

The ImageJ plugin NeuronJ was for neurite tracing and analysis. Each image consisted of a DAPI and MAP2 channel; in infected cultures, the GFP channel was referenced to determine areas of high infection. Prior to tracing, three regions of interest (ROI; 2000 x 2000 pixels) were overlayed onto each image. For ROIs without syncytia, five nuclei were chosen for neurite analysis. Neurites were traced from the soma to the longest branch. For ROIs containing syncytia, neurites were traced from the edge of the syncytia to the longest branch. (82–84)

### Viral Titers

To ensure adequate cell conditioning of all medium samples, collections occurred after a minimum of 48 hours in culture. Neuronal conditioned medium (nCM) was harvested at 7 dpi for both compound- and untreated neurons and subsequently was stored at −80C until use. To determine the viral titers of each CM sample, stocks were serially diluted, and the resulting dilutions applied to HCMV-naïve ARPE-19 cells. After 2 hours of exposure to undilute inoculum, fresh medium was added to dilute drug remaining in the conditioned medium. At 24 hours post infection (hpi), all inoculum-containing medium was removed from the cells and replaced with fresh medium. Cells were allowed to incubate in conditioned medium for 72 hours prior to being fixed and stained for HCMV Immediate Early Gene 1 (1°: mouse α-IE1, added 1:500, (Shenk lab) 2°: Goat α-mouse AF488 (1:1000)). As above, results are reported as infectious units per milliliter (IU/mL).

### Protein and DNA analyses

qRT-PCR was conducted to analyze relative amounts of various elements of the neuronal cytoskeleton. Using pelleted cells from 2, 4, 7, and 14 DPI, mRNA transcripts were isolated using the standard protocol for the RNeasy extraction kit (Qiagen). Isolated RNA was treated with RQ1 RNA-free DNAse (Promega) to ensure removal of contaminating genomic DNA prior to amplification. mRNA transcript amplification was conducted using random hexamers and the AMV Reverse Transcriptase System (Promega). Cytoskeletal elements were assessed using the following primer sets recommended by the PrimerBank tool (85, 86): TUBB3 (5′- ATCAGCAAGGTGCGTGAGGAG-3′ and 5′-TCGTTGTCGATGCAGTAGGTC-3′), MAP2 (5′- CGAAGCGCCAATGGATTCC-3′ and 5′-TGAACTATCCTTGCAGACACCT-3′), ACTB (5′- CATGTACGTTGCTATCCAGGC-3′ and 5′-CTCCTTAATGTCACGCACGAT-3′), NEFH (5′- CCGTCATCAGGCCGACATT-3′ and 5′-GTTTTCTGTAAGCGGCTATCTCT-3′), NEFM (5′- AGGCCCTGACAGCCATTAC-3′ and 5′-CTCTTCGGCTTGGTCTGACTT-3′), NEFL (5′- ATGAGTTCCTTCAGCTACGAGC-3′ and 5′-CTGGGCATCAACGATCCAGA-3′), MAPT (5′- CCAAGTGTGGCTCATTAGGCA-3′ and 5′-CCAATCTTCGACTGGACTCTGT-3′) and GAPDH (5′-GTGGACCTGACCTGCCGTCT-3′ and 5′-GGAGGAGTGGGTGTCGCTGT-3′) (Integrated DNA Technologies). Sequences were amplified using 2x SYBR Green Master Mix (Bio-Rad, Thermo Fisher). Data collection was accomplished using a Quantstudio 6 Flex real-time PCR machine (Thermo Fisher). Results for each gene were standardized against the expression of GAPDH transcripts.

Neurons intended for protein analysis were plated at a density of 5.0×10^6^ cells per well of a Geltrex-coated (Thermo Fisher) 6-well plate (5.2×10^4^ cells/cm^2^). At 51 days of differentiation, cells were infected with TB40/e-eGFP and allowed to persist for another 2-7 days post-infection. For 14 dpi samples, cells were grown for 84 days prior to infection. Collected cells were dislodged from their basement membrane using a P1000 pipette, pelleted by centrifugation (300 x g; 5 minutes), and frozen at −20°C until use. Cell pellets were treated with 50-150 µL of cold, pH=7.4 lysis buffer (150 mM NaCl, 50 mM Tris-HCl [pH 8.0], 1 mM EDTA, 1% Triton X-100 [v:v], 1% protease inhibitor [v:v], 1% phosphatase inhibitor [v:v]) for 20 minutes and lysed via sonication (2x 3s pulses; 30% amplitude). Sample concentrations were determined using the Pierce bicinchoninic acid (BCA) Assay Kit (Thermo Fisher) and a Glomax Microplate Reader (Promega). Utilizing 15-30 µg of protein (consistent within blots), sample contents were resolved by SDS-PAGE on a 4-20% acrylamide gradient gel (Bio-Rad). Separated protein bands were transferred to a polyvinylidene difluoride (PVDF) membrane (Millipore) using standard wet transfer procedure and a Mini TransBlot Cell (Bio-Rad). Using Intercept (PBS) Blocking Buffer (LI-COR), membranes were blotted to reduce the possibility of non-specific binding. Next, primary antibody solution (primary antibody, Intercept Blocking Buffer, 0.2% Tween-20 [v:v]) was applied, and membranes were incubated overnight at 4°C with agitation. Subsequently, membranes underwent three 5-minute washes with TBS-T (tris-buffered saline, 0.1% Tween-20) prior to applying secondary antibody solution (secondary antibody, Intercept Blocking Buffer, 0.2% Tween-20 [v:v], 0.02% sodium dodecyl sulfate [SDS; v:v]) for 30 minutes at room temperature. Following several washes (3x TBS-T, 1x TBS), blots were visualized using an Odyssey CLx fluorescent imaging system (LI-COR). Primary antibodies used in this study include: chicken anti-TUJ1 (1:250-500; GeneTex), chicken anti-MAP2 (1:500-1000; Invitrogen), mouse anti-MAP2 (1:1000, Invitrogen), mouse anti-NF68 (1:500, Sigma), rabbit anti-beta actin (1:500, Pierce), mouse anti-Tau46 (1:250-500, Cell Signaling Technologies), rabbit anti-TGN46 (1:500, Invitrogen), mouse anti-γ-tubulin (1:500-1000, Invitrogen), rabbit anti-GFP (1:1000, Invitrogen), mouse anti-IE1 (1:1000, Shenk Lab), and mouse anti-pp28 (1:400, Shenk Lab). Secondary antibodies used in this study include: donkey anti-rabbit 680RD (1:2000-3000, LI-COR), donkey anti-mouse 800CW (1:2000-3000, LI-COR), donkey anti-chicken 680RD (1:3000, LI-COR), and donkey anti-chicken 800CW (1:3000, LI-COR).

### Statistical Analysis

All statistical analyses contained within this study were completed using the GraphPad Prism software suite. Figure legends denote the specific statistical tests applied to each dataset, with significance being defined as * < 0.05 (** < 0.01, *** < 0.001, **** < 0.0001).

## ACKNOWLEDGEMENTS

The authors thank Drs. Felicia Goodrum and Eain Murphy for the fluorescently labeled viruses and Dr. Tom Shenk for the antibodies to HCMV IE1 and pp28. Further, we appreciate the help of all past and present Ebert and Terhune lab members for their helpful advice in the preparation of this manuscript. Thank you to Drs. Rebekah L Mokry and Suzette Rosas-Rogers for virus preparation and experimental consultation.

Funding support was provided by the National Institute of Allergy and Infectious Diseases division of the National Institutes of Health under award number R01AI132414 (S.S.T. and A.D.E.). All content herein is solely the responsibility of the authors and does not necessarily represent the official views of the National Institutes of Health. These studies have also been supported by a generous philanthropic gift by The Stead Family Foundation to define the impact of infection and inflammation on brain health.

Figures 1B, 2A, and 5A were designed using BioRender, per license agreement with the Medical College of Wisconsin. Licensing rights for these figure components are described in publication agreements: CU276HFCUI, RQ276HF8SE, and DC276HFQ36.

## AUTHOR CONTRIBUTIONS

Conceptualization: JWA, SST, ADE; data curation: JWA, ATS, KJP, JEG; data analysis and interpretation: JWA, ATS, KJP, JEG; funding acquisition: SST, ADE; resources: SST, ADE; supervision: SST, ADE; writing original draft, reviewing, and editing: JWA, ATS, KJP, JEG, SST, ADE. The authors declare no conflicts of interest.

## Notes

### Competing Interest Statement

The authors have declared no competing interest.

